# Genome-wide identification and transcriptional analyses of MATE transporter genes in root tips of wild *Cicer* spp. under aluminium stress

**DOI:** 10.1101/2020.04.27.063065

**Authors:** Xia Zhang, Brayden Weir, Hongru Wei, Zhiwei Deng, Xiaoqi Zhang, Yujuan Zhang, Xuexin Xu, Changxing Zhao, Jens D. Berger, Wendy Vance, Richard Bell, Yong Jia, Chengdao Li

## Abstract

Chickpea is an economically important legume crop with high nutritional value in human diets. Aluminium-toxicity poses a significant challenge for the yield improvement of this increasingly popular crop in acidic soils. The wild progenitors of chickpea may provide a more diverse gene pool for Al-tolerance in chickpea breeding. However, the genetic basis of Al-tolerance in chickpea and its wild relatives remains largely unknown. Here, we assessed the Al-tolerance of six selected wild *Cicer* accessions by measuring the root elongation in solution culture under control (0 µM Al^3+^) and Al-treatment (30 µM Al^3+^) conditions. Al-treatment significantly reduced the root elongation in all target lines compared to the control condition after 2-day’s growth. However, the relative reduction of root elongation in different lines varied greatly: 3 lines still retained significant root growth under Al-treatment, whilst another 2 lines displayed no root growth at all. We performed genome-wide identification of multidrug and toxic compound extrusion (MATE) encoding genes in the *Cicer* genome. A total of 56 annotated MATE genes were identified, which divided into 4 major phylogeny groups (G1-4). Four homologues to lupin *LaMATE* (> 50% aa identity; named *CaMATE1-4*) were clustered with previously characterised MATEs related to Al-tolerance in various other plants. qRT-PCR showed that *CaMATE2* transcription in root tips was significantly up-regulated upon Al-treatment in all target lines, whilst *CaMATE1* was up-regulated in all lines except Bari2_074 and Deste_064, which coincided with the lines displaying no root growth under Al-treatment. Transcriptional profiling in five *Cicer* tissues revealed that *CaMATE1* is specifically transcribed in the root tissue, further supporting its role in Al-detoxification in roots. This first identification of MATE-encoding genes associated with Al-tolerance in *Cicer* paves the ways for future functional characterization of MATE genes in *Cicer* spp., and to facilitate future design of gene-specific markers for Al-tolerant line selection in chickpea breeding programs.

## Introduction

Chickpea (*Cicer arietinum* L.) has become a valued grain legume worldwide, ranking second in area and third in production after soybean and pea (FAO, 2017). Chickpea seed is rich in protein, minerals, vitamins, and fibre, which provides many health benefits in diets ^1^, thus playing a critical role in human nutritional security. Over 60% of world chickpea production is from India, whilst Australia, Canada, and Argentina have seen increasing chickpea production in recent years, and have become leading chickpea exporters ^2^. During the past two decades, the world production of chickpea has increased steadily from ∼7 million tons to ∼14.5 million tons (FAO, 2019). However, chickpea yield has regained relatively stagnant.

Aluminium (Al) toxicity has been recognized as one of the major soil constraints for crop production. Around 30∼40% of the arable soils in the world are acid soils, and the area and severity continues to increase in due to factors such as acid rain, intensive agriculture, and the continued application of ammonium-based nitrogen fertilizers ^3^. The toxic Al^3+^ species significantly inhibits root elongation, thereby impairing nutrient and water uptake, and causes enormous crop yield loss. In chickpea, Al stress could cause inhibition of root growth, and possibly nodulation and nitrogen fixation also ^4,5^. In India ^6^ and Australia ^7^, both major chickpea producing countries, acidic soils account for a large proportion of the arable land. Thus, improved Al tolerance within chickpea cultivars would lead to higher crop yield on acid soils and the possibility of expanding chickpea production on soils where Al toxicity currently hampers cultivation.

Plants have developed various mechanisms to alleviate Al toxicity under acidic soils. The major mechanism is through the Al-activated release of organic acids from root tips ^8^. In barley, Al tolerance is achieved by the Al-induced secretion of citrate from barley roots, which chelates the toxic Al^3+^ in acidic soils ^9^. The secretion of citrate is facilitated by *HvAACT1* (Al-activated citrate transporter) gene encoding an enzyme in the multidrug and toxic compound extrusion (MATE) family ^9,10^. MATE transporters occur widely in nature, transporting substrates such as organic acids, plant hormones and secondary metabolites in both prokaryotes and eukaryotes ^11^. Homologous MATE proteins with similar citrate transport functions have been identified from wheat ^12^, maize ^13^, sorghum ^14^, rice ^15^, and Arabidopsis ^16^. In addition to the citrate transporter MATE, another Al-activated malate transporter (ALMT) has also been reported in many plants and is associated with the malate-mediated Al detoxification ^17,18^. Genetic studies on the Al-tolerance mechanism in grain legumes are still very limited. Several transcriptome analyses in root tips of legume plants indicated that MATE encoding genes are transcriptionally responsive to Al-treatment, and may have a similar Al-tolerance function ^19-21^.

In chickpea, the genetic basis of Al-tolerance remains obscure. Preliminary investigations have indicated that acid tolerance variations are present across different genotypes ^22,23^. Using two genotypes of varying Al-tolerance, Singh et al. ^24^ showed that Al-tolerance in chickpea may be controlled by a single dominant gene. However, no candidate gene has been identified to date. Furthermore, the current chickpea germplasm collection contains limited genetic variation related to biotic and abiotic stressors ^25,26^, which hinders the breeding progress for higher chickpea grain yield. The wild progenitor of chickpea (*Cicer reticulatum*) and its close relative, *C. echinospermum*, provide diverse gene pools for chickpea improvement that was recently widened by collection throughout SE Anatolia, Turkey where sampling covered a wide range of locations, climates and soil types ^27^. Interestingly, these two wild relatives of chickpea are found in different soil types: biologically derived limestone and sandstone soils for the former contrasting with geologically derived basaltic soils for the latter ^27^.Collection sites differ in terms of climate and soil properties: *C. reticulatum* collection site soils are more fertile and more alkaline than those where *C. echinospermum* was collected ^27^.Most importantly, *C. reticulatum* and *C. echinospermum* have no reproductive barrier with domesticated chickpea, therefore traits diversity in these wild *Cicer* spp. can be readily introduced in chickpea breeding programs ^26^.

In this study, we aim to explore the Al-tolerance variation within and between these two wild *Cicer* species, and identify the potential candidate genes contributing to Al-tolerance. Selected wild *Cicer* accessions were germinated and grown in a solution culture system under control and Al-treatment conditions. Al-tolerance was tested based on root elongation measurements ^23^. Genome-wide survey and phylogeny analyses of the MATE gene family in chickpea were performed. qRT-PCR experiments on two putative MATE candidate genes were carried out. This study is the first report of MATE-encoding genes transcriptionally associated with Al-tolerance in wild *Cicer* root tips, facilitating the future design of gene-specific markers for improved Al-tolerance in chickpea breeding programs.

## Results

### Effects of aluminium treatment on root growth

The resistance to Al toxicity was assessed by measuring the root elongation in solution culture under control (0 µM Al^3+^) and Al treatment (30 µM Al^3+^) conditions. We included six wild *Cicer* accessions displaying varying degrees of acid tolerance from a previous preliminary screening test ^28^. Under control condition (**Figure 1A**), the absolute root lengths ranged from 25 mm to 60 mm before and after 2-days (48 h) cultivation, reflecting the phenotypic variation among these chickpea lines. In particular, lines Bari2_074 and Deste_064 have relatively short root length (∼27 mm), whilst the other four lines have longer root lengths (> 35mm). Nested ANOVA showed significant differences both within and between species (**Supp. S1**). The highest root growth was observed in Karab_062 and Kayat_064 (*C. echinospermum* and *C. reticulatum*, respectively), followed by Sarik_073, CudiB_008B (both *C. reticulatum*) and Deste_064 (*C. echinospermum*). Significant 4-way interactions (P<0.001) indicate growth differences among *Cicer* lines across the 2 Al treatments. While 30 µM Al^3+^ reduced root extension in almost all varieties, the roots of Karab_062, Kayat_064, and Sarik_073 grow significantly longer over 2 days, whereas the remaining varieties are unable to do this (**Figure. 1B**). *C. arietinum* PBA HatTrick and *C. reticulatum* Bari2_074 were the exception, displaying no growth over time under both control and treatments (see **Supp. S1** for inter-line and inter-species statistical assessments).

**Figure 1.**
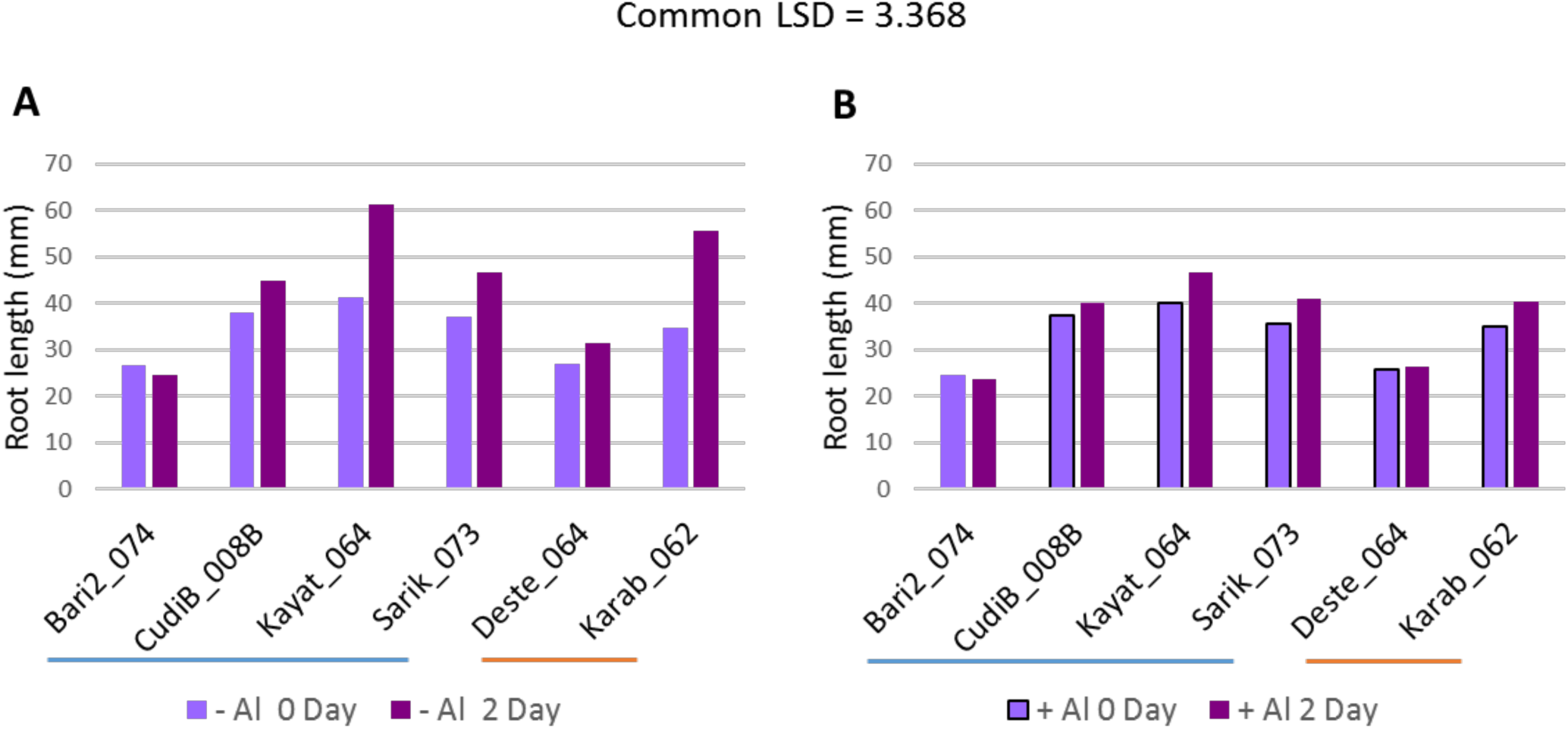
Root growth under control and Al treatment conditions for different wild *Cicer* accessions. Measurement of root length of *Cicer* seedlings before (0 Day) and after (2 Day) hydroponic cultivation in **A)** Control (– Al) and **B)** Aluminium treatment (+ Al) conditions. Al treatment contains 30 µMol Al^3+^; indicates *C. reticulatum* and indicates *C. echinospermum*. Least significant difference (LSD) = 3.368 for the 4-way ANOVA within variety (var), species (sp), Al-treatment (Al), and root growth (time). (see **Supp S1** for detailed statistics)

### Identification of candidate genes

The MATE gene is known to encode a citrate transporter which secretes citrate that detoxifies the free Al^3+^ in the rhizosphere in acidic soil. To identify the putative MATE transporter in the chickpea genome, the predicted amino acid sequences of the *Cicer* genome (NCBI BioProject: PRJNA190909) were searched using the MATE domain profile (Pfam ID: PF01554). A total of 56 unique peptide sequences containing the MATE domain were identified (**Supp. S2**). Several homologous MATE genes in lupin, soybean, Arabidopsis, barley and rice have been shown to play a critical role in Al resistance. To identify the orthologous MATE genes in *Cicer*, the amino acid sequence of lupin LaMATE (Uniprot ID: Q3T7F5) was used for the homology search against the *Cicer* genome. A total of 4 putative MATE homologues XP_004499881.1 (CaMATE1, 66.37% identity), XP_004510955.1 (CaMATE2, 60.93%), XP_004486970.1 (CaMATE3, 55.22%) and XP_004516070.1 (CaMATE4, 50.68%) were identified. The gene annotation of the homology search hits can be found in **Table 1**.

**Table 1.** List of homologous MATE gene hits in chickpea. Gene annotation was based on genome assembly ASM33114v1.

| Gene ID | Chromosome | Location | Protein ID | Annotation | aa Identity with LaMATE |
| --- | --- | --- | --- | --- | --- |
| LOC101509308<br>(CaMATE1) | Ca5 | NC_021164.1<br>(16522379..16529693) | XP_004499881.1 | protein DETOXIFICATION 43-like; MATE family; TIGR00797 | 66.37% |
| LOC101514527<br>(CaMATE2) | Ca7 | NC_021166.1<br>(39786461..39798690) | XP_004510955.1 | protein DETOXIFICATION 43-like; MATE family; TIGR00797 | 60.93% |
| LOC101497782<br>(CaMATE3) | Ca1 | NC_021160.1<br>(11155685..11160403) | XP_004486970.1 | protein DETOXIFICATION 42-like; MATE family; TIGR00797 | 55.22% |
| LOC101509930<br>(CaMATE4) | unknown | NW_004516700.1<br>(143529..149043) | XP_004516070.1 | protein DETOXIFICATION 42-like; MATE family; TIGR00797 | 50.68% |

### Phylogenetic analysis of MATE gene family

To investigate the evolutionary relationship of the identified MATE genes with their MATE homologues in the *Cicer* genome and other plants, a neighbour joining phylogeny was developed (**Figure 2**). Out of the 56 MATE transporters identified, 2 partial proteins were excluded from the phylogeny reconstruction. The developed phylogeny also included 31 previously studied MATE homologues from different plant species. As shown in **Figure 2**, *Cicer* MATE proteins divided into 4 major phylogenetic groups G1-4. The target MATE proteins XP_004499881.1 (CaMATE1) and XP_004486970.1 (CaMATE2) identified in the present study are present in group G4, which also contained soybean GmFRD3b ^29^, lupin LaMATE ^30^, Arabidposis AtFRD3 ^31^, and *Eucalyptus* EcMATE1 ^32^ and the other characterised MATE genes related to aluminium resistance in monocot plants, supporting the potential function of these two *Cicer* MATE genes in Al detoxification. Within group G4, CaMATE1 and CaMATE2 were clustered with other legume MATE homologues, GmFRD3b and LaMATE. Compared to CaMATE2, CaMATE1 seems to display a relatively closer relationship with GmFRD3b. Another two *Cicer* MATEs XP_004486970.1 and XP_004516070.1 were present in a separate subgroup with AtMATE and cabbage BoMATE. Interestingly, this subgroup tends to have a closer relationship with the monocot orthologues than CaMATE1 and CaMATE2.

**Figure 2.**
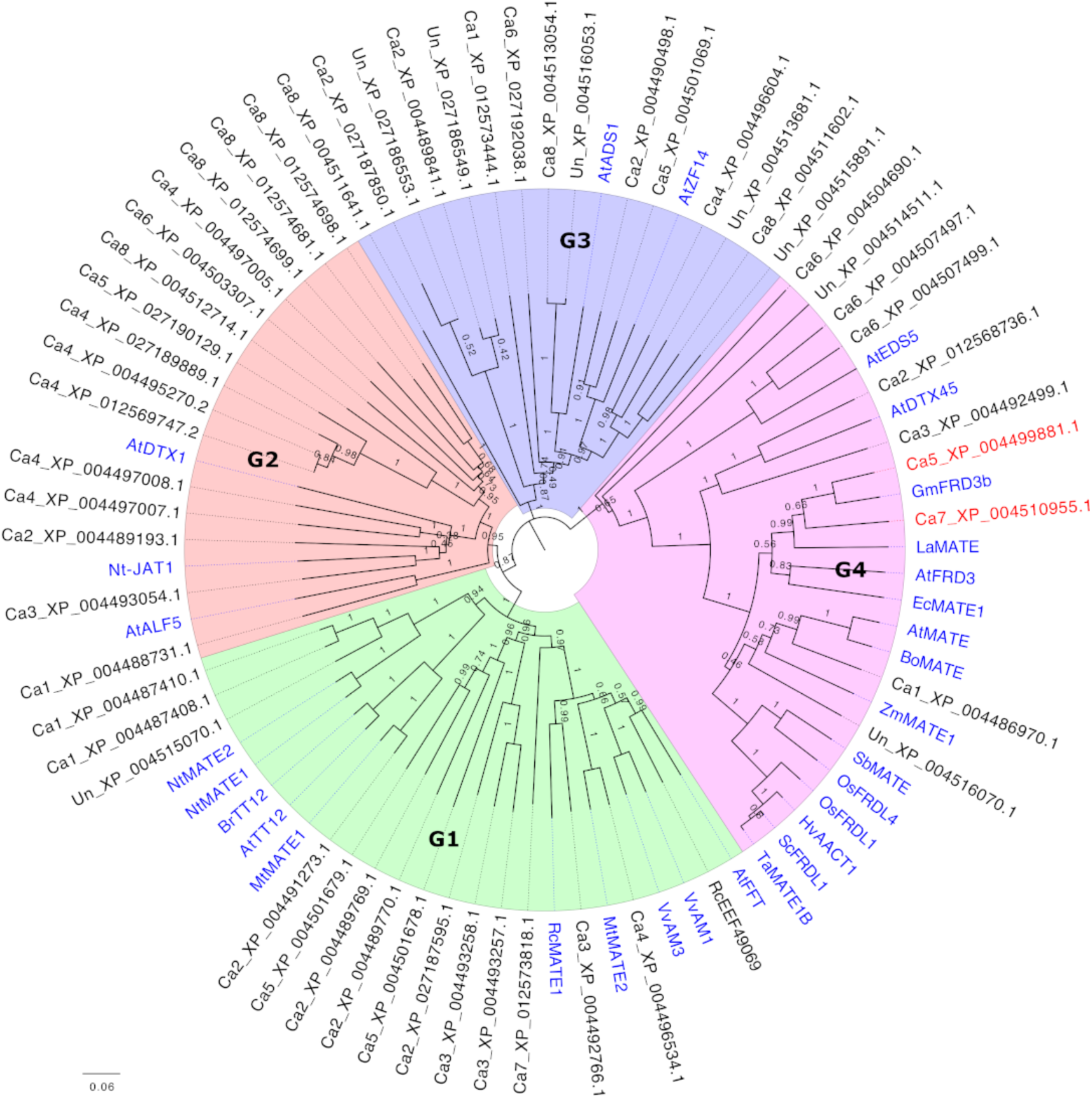
Phylogeny of MATE homologous genes in chickpea and other plants. The NJ phylogeny includes the chickpea protein sequences containing the MatE (PF01540) domain (retrieved from NCBI database BioProject: PRJNA190909). Previously characterised homologous MATE proteins were included as references (highlighted in blue). The target MATE, CaMATE1 and CaMATE2, were in red. The Bootstrap support (1000 times iteration) was indicated above each branch.

### Synteny and gene structural analyses

Depending on the different genetic mechanisms, gene family expansion can be attributed to four gene duplication types: whole genome duplication (WGD)/segmental duplication, tandem duplication, proximal duplication and dispersed duplication. To investigate the evolutionary origin of the *Cicer* MATE gene family, synteny and gene structural features were analysed based on the developed phylogeny. As shown in **Figure 3**, a total of 6 collinear gene pairs and 11 tandem gene pairs (covering 27 genes) were identified within the *Cicer* MATE family, suggesting these genes have originated from WGD/segmental duplication and tandem duplication, respectively. These two types of duplication account for almost half of the *Cicer* MATE genes, whilst the other genes were classified as dispersed or proximal duplication, which include *CaMATE1-4*. Gene structural analyses showed that G1 and G2 MATE genes generally have similar exon-intron profiles, suggesting these two groups may have originated from a recent divergence event. In contrast, G3 and G4 displayed distinct gene structural profiles from G1 and G2. In particular, *CaMATE1* and *CaMATE2* contained 12 exons, whilst *CaMATE3* and *CaMATE4* had 13 exons, which is consistent with their phylogeny relationship.

**Figure 3.**
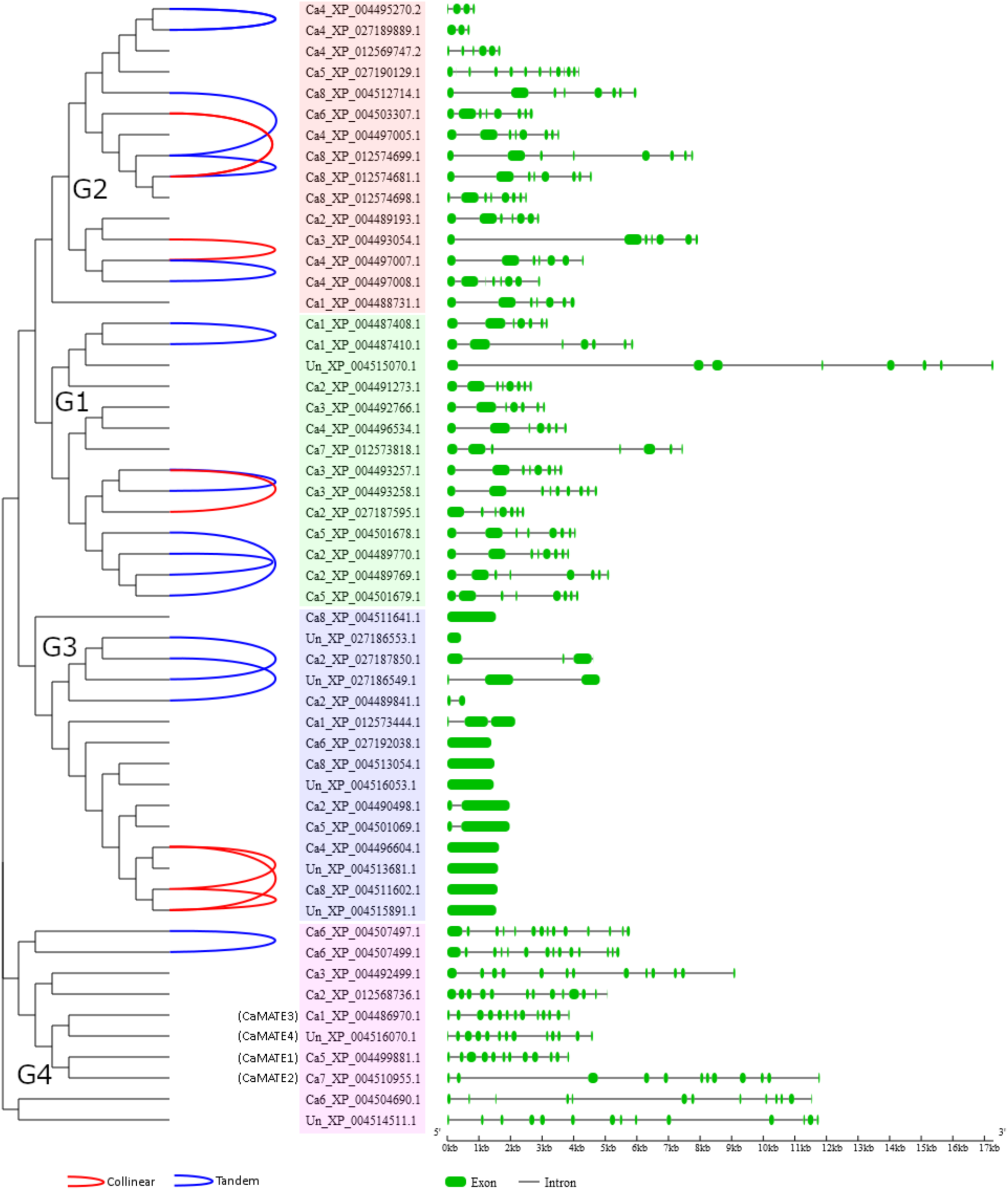
Synteny and gene structural analyses of chickpea MATE family. The synteny and gene structural features were displayed based on the developed MATE phylogeny. On the left, identified collinear and tandem duplication gene pairs were linked by red and blue lines, respectively. In the middle, phylogeny groups G1-G4 were highlighted in pink, blue, light green, and brick red, respectively. On the right, exon and intron features were displayed in green rectangle and black line, respectively.

### qRT-PCR analyses

The MATE family genes encode transporter proteins that transport organic acid molecules, such as citrate or malate, from root to soil, thus facilitating the chelation of the toxic Al ions. The most active tissue in which the MATE genes are highly transcribed is the root tip ^11^.

To validate the potential function of *Cicer* MATE genes in Al tolerance, the expression levels of the MATE genes in root tips (1-2 cm) and their response to Al treatment were investigated. Two representative MATE genes, *CaMATE1* and *CaMATE2* that are most closely related to the previously characterized *AtFDR3* and *AtMATE*, were selected for qRT-PCR experiments. Under the control condition (0 Al^3+^), the transcription level of *CaMATE1* varied greatly across the six wild Cicer accessions, with line Deste_064 displaying the highest expression, followed by line Bari2_074, and then by Line CudiB_008B (**Figure 4A**), whilst chickpea lines Kayat_064, Sarik_073 and Karab_062 demonstrated the lowest and similar expression of *CaMATE1*, which was less than a quarter of that in line Deste_064. After applying the Al treatment, the transcription of *CaMATE1* increased significantly in lines Kayat_064, Sarik_073 and Karab_062 by around ∼3 times. Moderate increase (∼1.5 times) of *CaMATE1* expression was observed in line CudiB_008B. In contrast, the transcription of *CaMATE1* under Al treatment decreased dramatically in line Deste_064 (∼0.4 times), and dropped slightly in line 59809. Notably, the downregulation of *CaMATE1* in lines Bari2_074 and Deste_064 coincided with the relatively short root length observed for these two chickpea lines. This also corresponds well with the observation that no root elongation was detected for these two lines after 4 days cultivation under both conditions.

**Figure 4.**
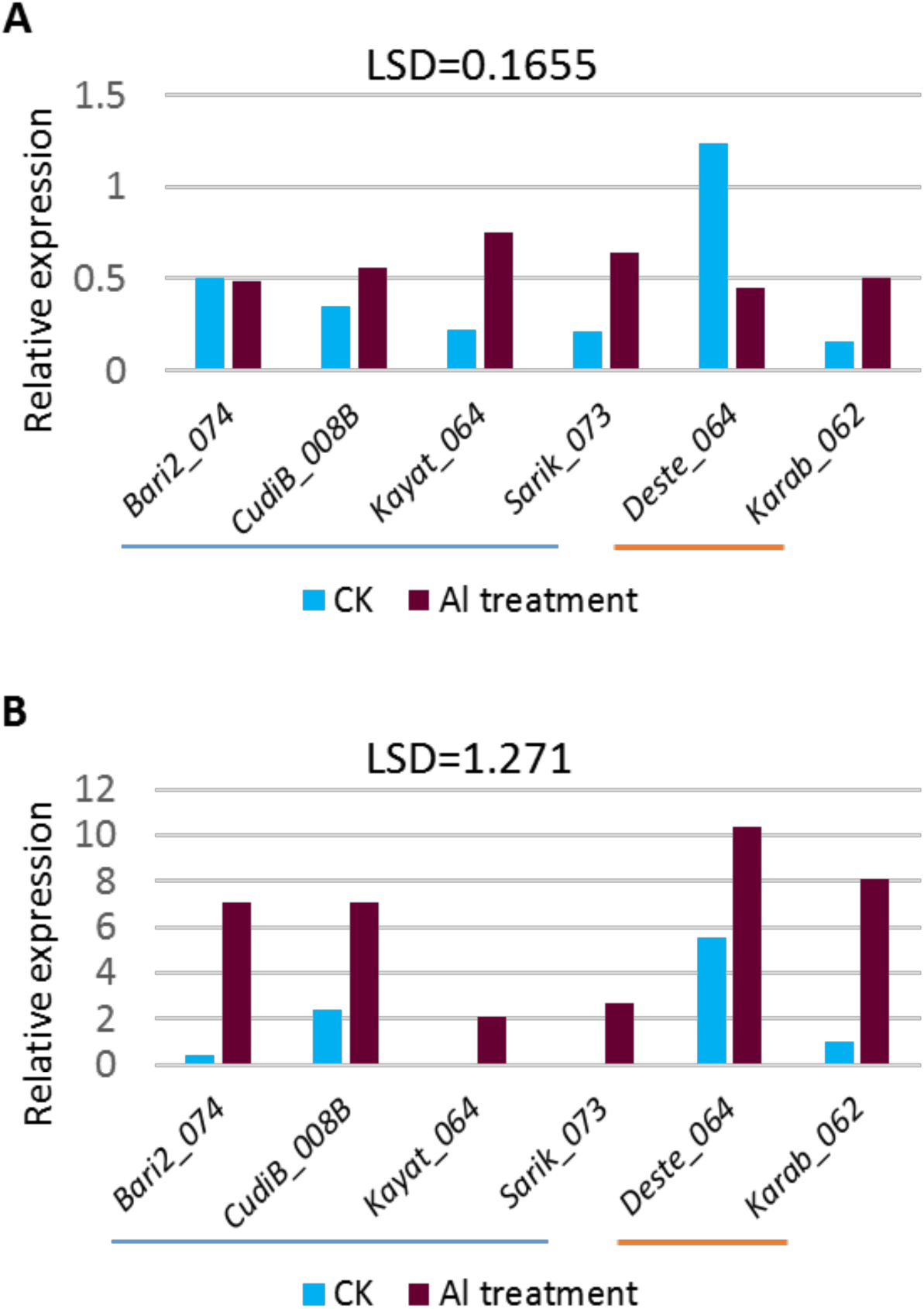
qRT-PCR analyses on *Cicer* MATE genes in root tips. The relative transcription of *CaMATE1* (**A**) and *CaMATE2* (**B**) was determined in six *Cicer* lines Bari2_074, CudiB_008B, Kayat_064, Sarik_073, Deste_064 and Karab_062 after 2 days hydroponic cultivation. The previously determined *CaCAC* was used as the reference gene. indicates *C. reticulatum* and indicates *C. echinospermum*. LSD values of 3-way ANOVA within Al-treatment (Al), varieties (var) and species (sp) for *CaMATE1* and *CaMATE2* are 0.1655 and 1.271, respectively. (see **Supp S1** for detailed statistics)

Similar to *CaMATE1*, the transcription of *CaMATE2* (**Figure 4B**) also differed greatly across different *Cicer* accessions. Under the control condition, the highest *CaMATE2* expression was detected in line Deste_064, followed by line CudiB_008B. However, the other lines had relatively low or barely any transcription of *CaMATE2*. Compared to the control condition, the Al treatment led to significant upregulation of *CaMATE2* in all *Cicer* lines studied. Under Al treatment, lines Deste_064, Karab_062, CudiB_008B and Bari2_074 displayed abundant *CaMATE2* transcription, which was approximately 3∼5 times that in lines Kayat_064 and Sarik_073.

### Transcriptome analyses

To further character the transcriptional profile of *Cicer* MATE genes, the transcriptional data of MATE-encoding genes in 5 different tissues (shoot, root, mature leaf, flower bud and young pod) were retrieved from the public database. As shown in **Figure 5**, 35 out of the 56 MATE genes in *Cicer* genome were identified with available transcriptional data. Based on the phylogeny clustering pattern, G1 and G4 genes tend to be expressed relatively higher in root tissues than G2 and G3, thus highlighting their potential involvement in Al tolerance. In contrast, most of G3 genes are barely transcribed in any of the 5 tissues studied, with the exception of Ca8_XP_004511641.1 which was moderately expressed in root. In addition, several MATE genes displayed a clear tissue-specific expression pattern, which include Ca2_XP_004491273.1 (G2) in young pod, Ca5_XP_004501069.1 (G3) in shoot, and Ca5_XP_004499881.1 (G4) in root. In particular, Ca5_XP_004499881.1, corresponds to CaMATE1 in the present study. The root-specific expression of CaMATE1 corroborates its proposed role in Al tolerance.

**Figure 5.**
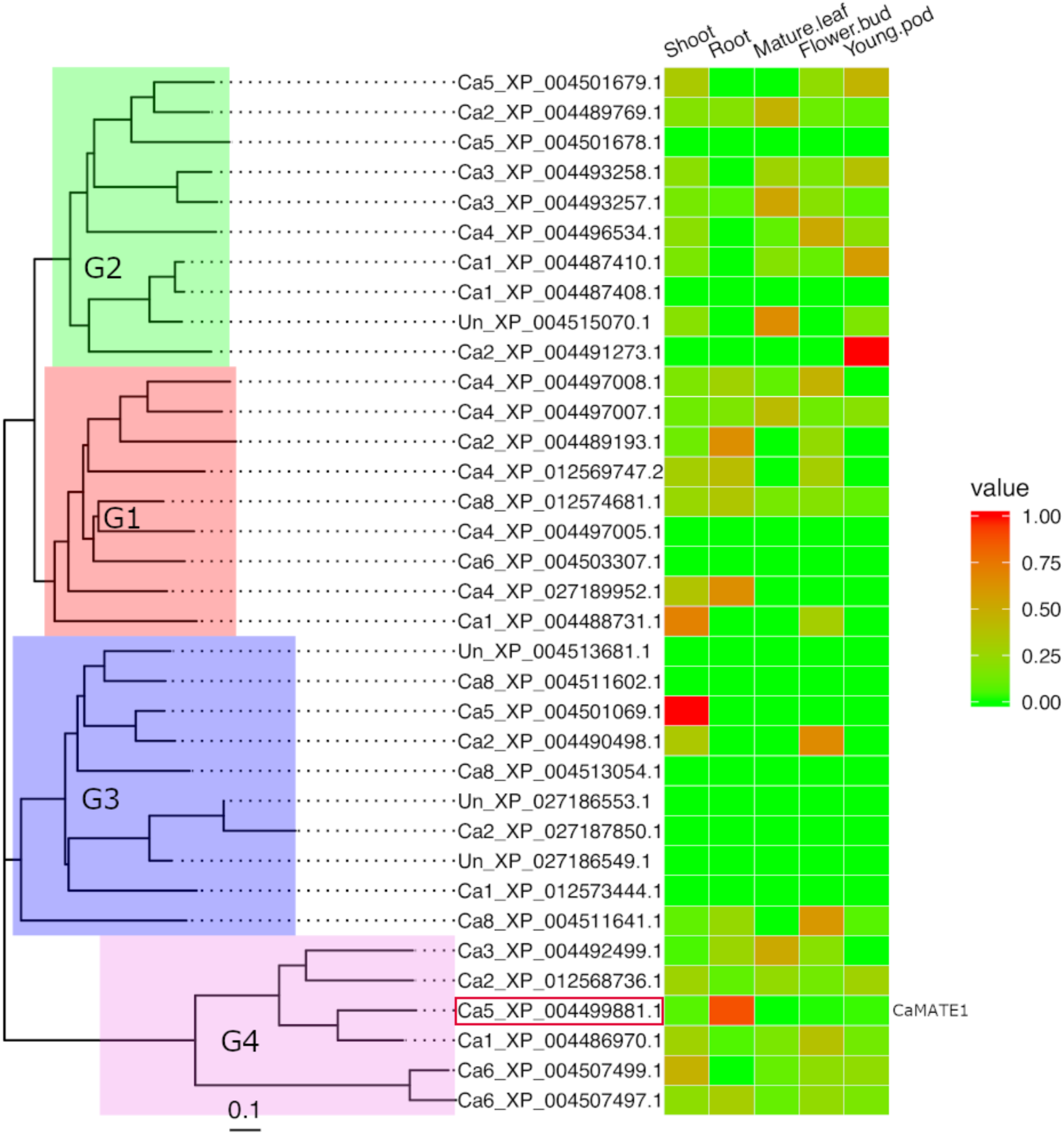
Transcriptional heat-map of *Cicer* MATE genes across different tissues. Transcriptional data for MATE domain containing genes in 5 different tissues (shoot, root, mature leaf, flower bud and young pod) were retrieved from chickpea transcriptome database (CTDB) and normalized based on individual genes. The normalized data were plotted in heatmap according to the clustering pattern (G1:green, G2:red, G3:blue, G4:pink) of an un-rooted neighbour-joining tree. The position of CaMATE1 (Ca5_XP_004499881.1) was highlighted in the red box.ed text.

## Discussion

Our results showed that there was significant variation in Al-tolerance among the target wild *Cicer* lines, thereby supporting the potential use of wild *Cicer* for Al-tolerance improvement in chickpea breeding. Chickpea is susceptible to Al-stress ^4,5^. To date, two studies have attempted to examine the genotypic variations against Al-stress. The assessment of Al-tolerance in 35 and 24 cultivated chickpea genotypes, respectively, have allowed the identification of relatively tolerant and sensitive chickpea lines ^23,24^. These Al-tolerant lines may be used for yield improvement in chickpea breeding. However, compared to the other crops species, the genetic diversity of chickpea germplasm against various other abiotic and biotic stresses is also relative narrow ^26,33^, which hinders the progress on chickpea breeding toward higher yield under unfavourable environmental conditions. The lack of sufficient genetic diversity in chickpea, however, can be complemented by some of its wild progenitors such as *C. reticulatum* and *Cicer echinospermum*, which displays no reproductive barrier with cultivated chickpea ^26,34^. Based on these observations, the current study attempted to evaluate the Al-tolerance variation in these two species. Our study is the first reported evaluation of Al-tolerance in wild *Cicer*.

The genetic basis of Al-tolerance in chickpea remains largely unknown. Based on the assessment of Al-sensitivity in the progeny of two chickpea parental lines, Singh et al. ^24^ determined that the Al-tolerance variation in the two parental lines may be controlled by a single dominant gene. However, the underlying candidate gene and its physiological mechanism were not identified. The Al-activated MATE transporter facilitates the secretion of citrate from the root apex, which is the major mechanism of Al-tolerance in many plants ^8^. The availability of the chickpea genomic data ^35^ has enabled the genome-wide survey of putative MATE-encoding genes in the present study. Based on the most recent chickpea genome annotation, we identified a total of 56 MATE homologues in *Cicer*, which is close to the 71 reported for Populus ^36^ but significantly less than the 117 for soybean ^21^. Phylogeny analysis suggested that the MATE gene family could be divided into four major subclades, which is similar with the observation made in other species such as soybean ^21^ and Populus ^36^. In our phylogeny, two *Cicer* MATE homologues were clustered each with the previously identified AtMATE and AtFRD3, respectively, which resembled the observation in Populus ^36^. In contrast, the soybean reference genome contained 4 close homologues each for AtMATE and AtFRD3 ^21^, which may result from its recent polyploidy.

Both *CaMATE1* and *CaMATE2*, representing the direct homologues to *AtFRD3*, were significantly upregulated upon Al-treatment. This observation is similar with the transcriptional upregulation for soybean *GmMATE75* ^21^, barley *HvAACT1* ^10^, and the Populus *PtrMATE1, PrtMATE2, PtrDXT2*, and *PtrDXT27* upon Al-treatment ^36^. In barley, Al-tolerant varieties displayed significantly longer root elongation that Al-sensitive lines, which is associated with higher HvAACT1 transcription in the root tips ^9^. In this study, we found that the transcriptional level of *CaMATE1* and *CaMATE2* were positively correlated to the genotypic variation in root elongation in wild *Cicer*. In particular, transcriptome profiling suggested that the transcription of *CaMATE1* may be root-specific, further supporting its role in Al-resistance. Thus, it would be intriguing for further study to verify if *CaMATE1* and *CaMATE2* may correspond to the monogenic Al-tolerance locus identified by Singh et al ^24^. In addition, future study can also be devoted to identify the genetic polymorphism of *CaMATE1* and *CaMATE2* across a much larger collection of chickpea and wild *Cicer* germplasm. Novel allele(s) associated with high Al-tolerance may be identified and used for chickpea breeding.

Generally, Al-tolerance in plants is a complex trait involving multiple gene families and pathways. In addition to the MATE-encoding genes, candidate genes from other pathways may also contribute to Al-tolerance in chickpea. For example, comprehensive transcriptome profiling in medicago and soybean root tips have revealed that many genes related to oxidative stress, transcriptional regulation, cell wall process, lignin deposition are also responsive to Al-treatment ^19,20^. Comparative transcriptome study is also necessary to unravel other potential genetic mechanisms associated with Al-tolerance in chickpea. Transgenic over-expression of ALMT homologues in medicago and soybean have also been shown to increase Al-resistance ^37,38^. In *Arabidopsis*, AtSTOP1, a C2H2 zinc finger transcription factor that regulates the expression of AtMATE and AtMLT1, is also involved in Al-tolerance ^39^.The AtSTOP orthologue in rice, OsART1, has also been characterized to be related to Al-tolerance ^40^. Recently, the effect of microRNAs on Al-tolerance in barley was tentatively investigated, providing new insights into this complex biological process ^41^. Therefore, it is necessary for future study to verify if a similar genetic basis for controlling Al-tolerance may be present in chickpea or not. On another note, legume plants including chickpea can characteristically form nodules in the root for N-fixation. Al in acidic soils may pose a constraint on nodule-formation due to its lethal effect on rhizobia ^42^. Therefore, for the improvement in chickpea production in acidic soil, attention should also be given to the rhizobia acidity tolerance. As an earlier study has shown, most acid-tolerant chickpea mesorhizobia showed transcriptional induction of major chaperone genes upon acid treatment, whilst the sensitive strains showed repression ^43^.

## Conclusions

The wild progenitors of chickpea provide a diverse gene pool for Al-tolerance in chickpea breeding. We assessed and verified the presence of significant Al-tolerance variation across 6 different wild *Cicer* genotypes. A genome-wide survey identified a total of 56 putative MATE-encoding genes in the chickpea genome. Results of phylogeny and transcriptional analyses revealed the positive role of *CaMATE1* and *CaMATE2* in Al-resistance in *Cicer* roots, and support their potential use in future chickpea breeding for yield improvement.

## Methods

### Plant materials, hydroponic cultivation and tissue sampling

A total of six wild *Cicer* lines, *C. reticulatum*: Bari2_074, CudiB_008B, Kayat_064, and Sarik_073; *C. echinospermum*: Deste_064 and Karab_062, were obtained from the germplasm collected from southeastern Anatolia, Turkey. Around 50 seeds for each line were used. Sterilised seeds (3% sodium hypochlorite for 5 minutes, followed by rinsing 5 times with de-ionized water) were placed on a petri-dish covered with wet paper towel to allow germination for 4 days.

On day 5, seedlings were transferred to 5-litre containers containing solution with constant aeration. All seedlings were initially in the control condition. The complete nutrient solution at pH 4.2 contained (μM): CaCl_2_.2H_2_O, 400; KNO_3_, 650; MgCl_2_.6H_2_O, 250; (NH_4_)_2_SO_4_, 10; NH_4_NO_3_, 40; H_3_BO_3_, 23; MnCl_2_.4H_2_0, 9; Na_2_MoO_4_.2H_2_O, 0.1; ZnSO_4_.7H_2_O, 0.8; CuSO_4_.5H_2_O, 0.3; Na_2_HPO_4_, 5. Iron (20 µM) was supplied as Fe-EDTA prepared from equimolar amounts of FeCl_3_.6H_2_O and Na_2_EDTA at pH 4.2. On day 6 the root length was measured using a vernier caliper before returning seedlings to the solution containers with either control (pH 4.27) or the Al-treatment solution (pH 4.25) which contained 30 µMol Al3+ added as AlCl_3_.6H_2_O.

After 48 hours in treatment solutions, the root length was measured again. The root tips (1-2 cm) were sampled using a scalpel blade, snap-frozen in liquid nitrogen, and stored in −80 °C until RNA extraction. Three biological replicates were included for each line, with each replicate comprising 5 seedlings.

### Sequence retrieval and primer design

The amino acid sequence of lupin LaMATE was used to blastp against the NCBI chickpea genome data (BioProject: PRJNA190909). The genomic DNA sequence and transcript sequence for the target MATE genes were retrieved. qRT-PCR primers spanning the introns were designed using the RealTime PCR Design Tool (Integrated DNA Technologies, US, https://sg.idtdna.com/scitools/Applications/RealTimePCR/)

### Phylogeny development

The predicted amino acid sequences for the chickpea genome were downloaded from the NCBI database (BioProject: PRJNA190909). The MATE domain profile file (MatE.hmm) was downloaded from the Pfam database (https://pfam.xfam.org/). The hmmscan program (http://hmmer.org/) was used to identify the sequences containing the MATE domain. The amino acid sequences of previously reported MATE proteins were retrieved from the Uniprot database (https://www.uniprot.org/). A list of previously characterized MATEs was retrieved from a recent study ^36^. For phylogeny inference, amino acid sequence alignment was performed using MUSCLE (8 iterations) ^44^. Phylogeny was developed using the Neighbour Joining (NJ) method implemented in MEGA 7.0 ^45^ with the p-distance substitution model. 1000 times bootstrap support was calculated for the developed NJ tree. Tree annotation was performed using the FigTree tool at http://tree.bio.ed.ac.uk/software/figtree/.

### Synteny and gene structural analyses

Synteny and gene duplication pattern were analysed using MCScanX software ^46^. Chickpea genome annotation data were downloaded from the NCBI database (BioProject: PRJNA190909). Intra- and inter-species genome comparisons were performed using the standalone NCBI-BLAST-2.2.29 tool with an E-value threshold of 1e-05, restricting the maximum hit number to 5. Collinear and tandem gene pairs were displayed using the family_tree_plotter tool in MCScanX package ^46^. Gene structure features were displayed using the GSDS 2.0 tool ^47^.

### RNA extraction and cDNA synthesis

The frozen root tips samples were ground into a fine powder using a pestle and a mortar pre-cooled in liquid nitrogen. RNA extraction was carried out using Trisure® (Bioline, Australia) by following the manufacturer’s instruction. ∼100 mg of ground tissue was used for each extraction. cDNA library construction was performed using SensiFAST™ cDNA Synthesis Kit (Bioline, Australia).

### RT-qPCR

The RT-qPCR experiments were carried out using SensiFAST™ SYBR No-ROX Kit (Bioline, Australia). Each reaction contains 5 µl SensiFAST mix, 4.2 µl cDNA template, 0.8 µl forward/reverse primers (500 nM). The RT-PCR primers are forward: CCTGCAGTGCTTCTCTCTTT & reverse: GCATACCCGGAAACTATGACA for CaMATE1 and forward: GGCTTCCTTCAAGCTTCAATTC & reverse: GCAGGAGCACCAAATGATCTA for CaMATE2. RT-qPCR reaction was performed using the ViiA7 Real-Time PCR System (Thermo Fisher, USA) in 384-well plates. The previously tested chickpea CaCAC gene was used as a reference gene ^48^. Three replicates were included for each sample. Each sample was run in three technical replicates. The transcription values were calculated using the comparative Ct method (2^-ΔCt^) ^49^.

### Transcriptional data mining

The transcriptional data of MATE-encoding genes were retrieved from the chickpea transcriptome database (CTDB) at http://www.nipgr.ac.in/ctdb.html. The obtained transcriptional data in RPKM unit was normalized based on individual gene in different tissues. A separate un-rooted neighbour-joining phylogeny was developed using MEGA7.0 ^45^, which covers MATE genes with transcriptional data available. The transcriptional heat-map data was plotted using the ggtree ^50^ R package based on the phylogeny clustering pattern.

### Statistics analysis

Linear regression and ANOVA routines in Genstat V20 (VSN International, Hemel Hempstead, UK) were used to analyse the data, using residual plots to check for normality and identify outliers. Varieties were nested within species, using Al treatment and time as factors, except in linear regression, where time was treated as a variate.

## Acknowledgement and Funding

The authors would like to acknowledge the chickpea research community for making the genomic and transcriptome data available to the public. The work was funded through GRDC project UMU00044. Genetic material was made available through the collection program of GRDC project CSP00185. Seeds were made available through SMTA with the Australian Grains Gene bank under agreement with AARI (Izmir Genebank).

## Author contribution

CL, RB & WV supervised the study. XZ, WV, HW & ZD performed hydroponic tests. BW, XZ, HW & ZD did qRT-PCR experiments under YJ’s supervision. JDB provided chickpea materials and data analysis. XQZ, XX & CZ assisted laboratory experiments. YJ&YZ performed bioinformatics and data analyses. YJ wrote the manuscript. JDB, RB, CL & WV provided valuable revisions. All authors have read the manuscript.

## Supplementary materials

**Supplementary file S1.**
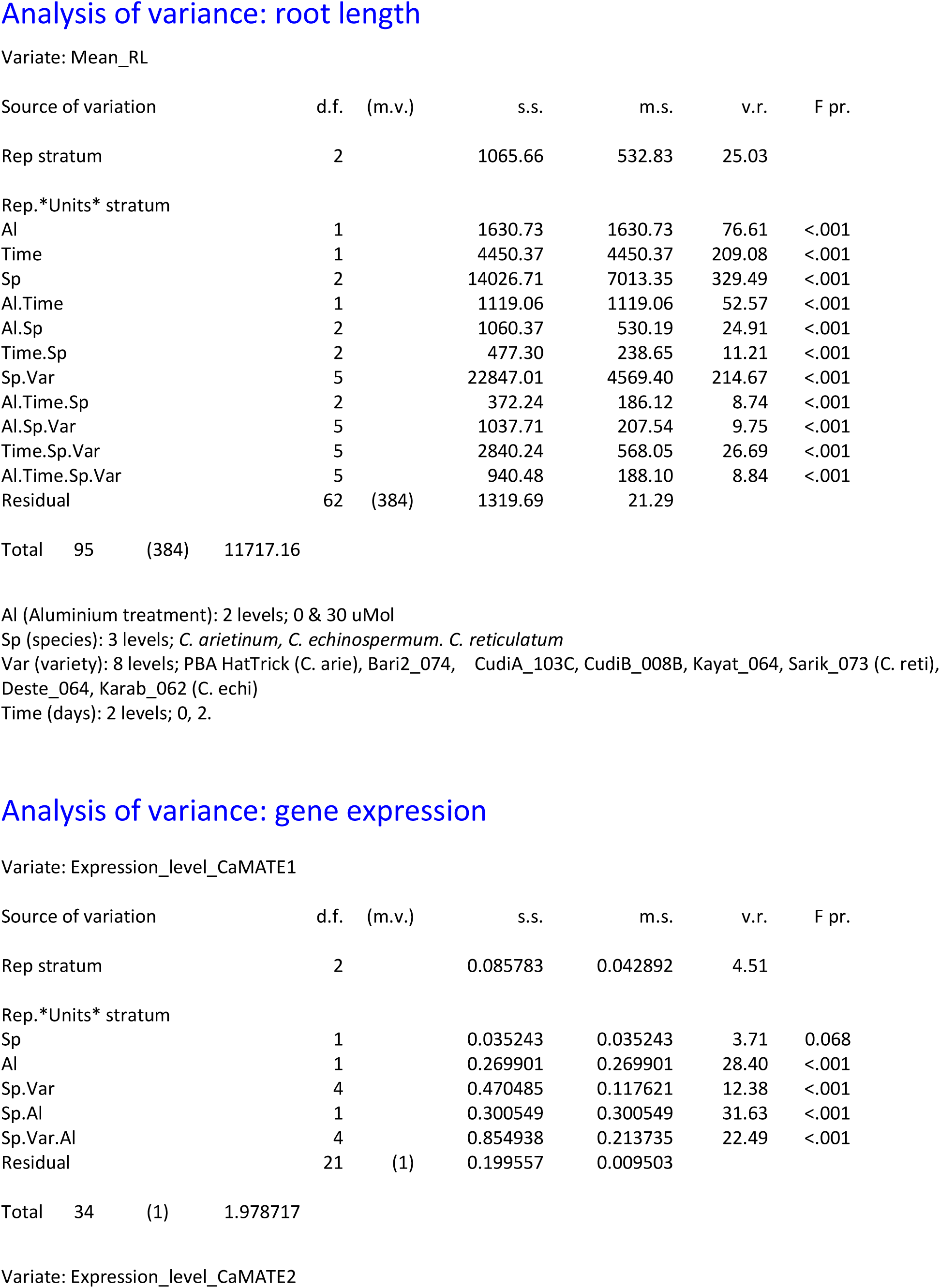
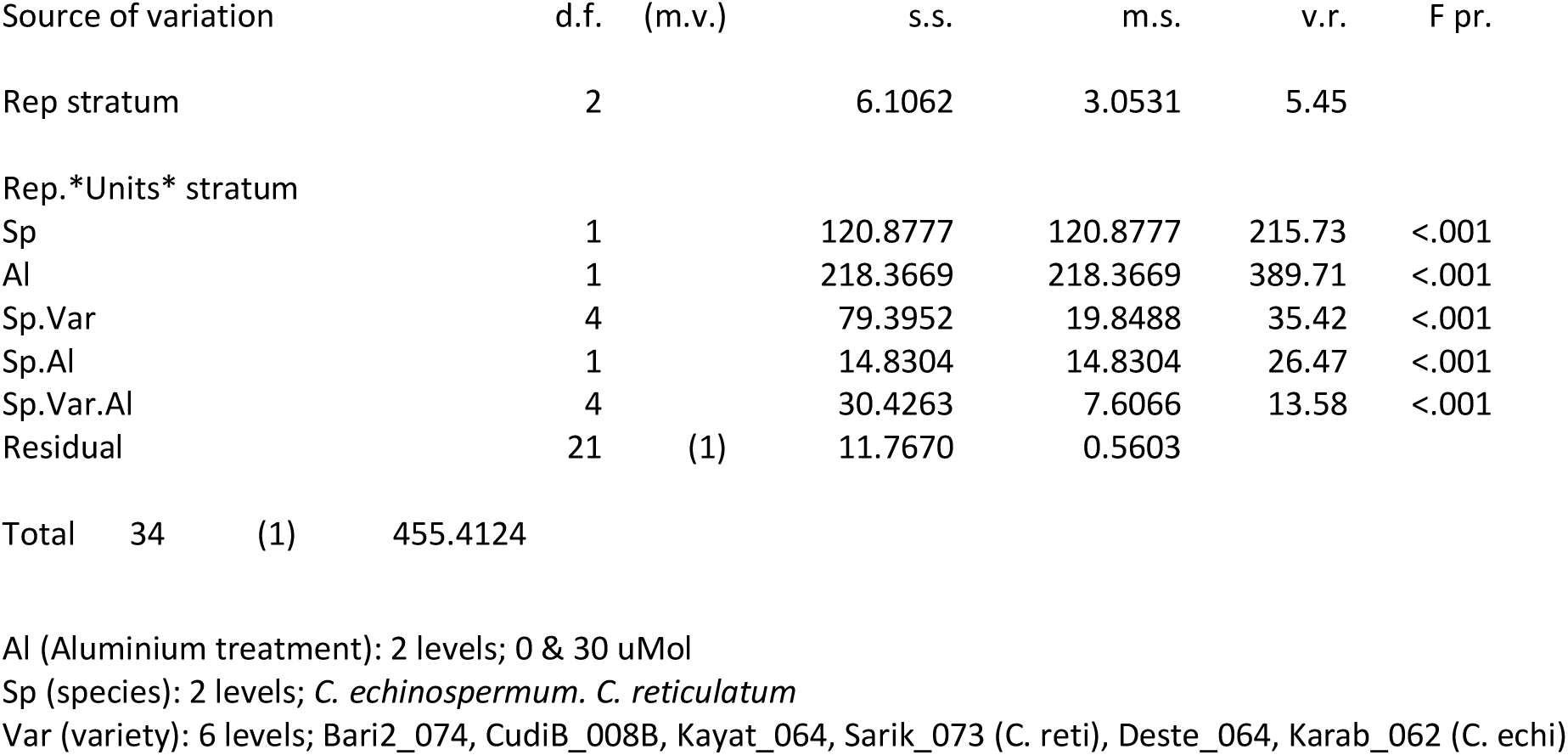
Inter-line and inter-species variance tests on the root growth of wild Cicer lines.

**Supplementary file S2.** List of identified putative MATE encoding genes in chickpea.

| Protein ID | Gene ID | Chr | Start | End | Strand |
| --- | --- | --- | --- | --- | --- |
| XP_004486970.1 | LOC101497782 | Ca1 | 11156189 | 11160059 | + |
| XP_004487408.1 | LOC101506667 | Ca1 | 14022099 | 14025275 | - |
| XP_004487410.1 | LOC101507327 | Ca1 | 14028317 | 14034199 | - |
| XP_012573444.1 | LOC101508596 | Ca1 | 17930045 | 17932194 | + |
| XP_004488731.1 | LOC101496368 | Ca1 | 41825183 | 41829212 | - |
| XP_004489193.1 | LOC101502073 | Ca2 | 1657967 | 1660872 | + |
| XP_027187595.1 | LOC101495061 | Ca2 | 5907697 | 5910130 | - |
| XP_004489769.1 | LOC101503133 | Ca2 | 7678614 | 7683731 | + |
| XP_004489770.1 | LOC101503455 | Ca2 | 7699482 | 7703334 | + |
| XP_027187850.1 | LOC113785478 | Ca2 | 8425149 | 8429765 | + |
| XP_004489841.1 | LOC101500282 | Ca2 | 8440064 | 8440631 | + |
| XP_004490498.1 | LOC101495941 | Ca2 | 24037644 | 24039619 | - |
| XP_004491273.1 | LOC101513987 | Ca2 | 33782028 | 33784704 | + |
| XP_012568736.1 | LOC101497360 | Ca2 | 36396414 | 36401491 | + |
| XP_004492499.1 | LOC101505307 | Ca3 | 20176048 | 20185161 | + |
| XP_004492766.1 | LOC101513768 | Ca3 | 23326545 | 23329636 | - |
| XP_004493054.1 | LOC101498359 | Ca3 | 25203613 | 25211545 | - |
| XP_004493257.1 | LOC101509490 | Ca3 | 27417502 | 27421139 | + |
| XP_004493258.1 | LOC101509807 | Ca3 | 27431809 | 27436550 | + |
| XP_012569747.2 | LOC101498383 | Ca4 | 336802 | 338479 | - |
| XP_004495270.2 | LOC101498724 | Ca4 | 347598 | 348469 | - |
| XP_027189886.1 | LOC113786338 | Ca4 | 349135 | 350104 | - |
| XP_027189889.1 | LOC105851853 | Ca4 | 367668 | 368375 | - |
| XP_027189952.1 | LOC113786368 | Ca4 | 369084 | 370001 | - |
| XP_004496534.1 | LOC101514779 | Ca4 | 12210225 | 12214002 | + |
| XP_004496604.1 | LOC101509286 | Ca4 | 12821840 | 12823477 | - |
| XP_004497005.1 | LOC101500523 | Ca4 | 15378577 | 15382117 | - |
| XP_004497007.1 | LOC101501055 | Ca4 | 15386018 | 15390333 | + |
| XP_004497008.1 | LOC101501367 | Ca4 | 15392608 | 15395544 | + |
| XP_027190129.1 | LOC101505782 | Ca5 | 288578 | 292755 | - |
| XP_004499881.1 | LOC101509308 | Ca5 | 16525537 | 16529392 | + |
| XP_004501069.1 | LOC101492659 | Ca5 | 32893590 | 32895564 | - |
| XP_004501678.1 | LOC101494285 | Ca5 | 37657645 | 37661708 | + |
| XP_004501679.1 | LOC101494597 | Ca5 | 37664214 | 37668363 | + |
| XP_004503307.1 | LOC101506763 | Ca6 | 2418208 | 2420913 | - |
| XP_004504690.1 | LOC101492232 | Ca6 | 13938414 | 13949963 | - |
| XP_027192038.1 | LOC101511466 | Ca6 | 37038724 | 37040118 | - |
| XP_004507497.1 | LOC101494100 | Ca6 | 58728279 | 58734058 | - |
| XP_004507499.1 | LOC101494621 | Ca6 | 58743260 | 58748708 | - |
| XP_012573818.1 | LOC101512584 | Ca7 | 18527103 | 18534560 | + |
| XP_004510955.1 | LOC101514527 | Ca7 | 39786668 | 39798457 | - |
| XP_004511602.1 | LOC101504565 | Ca8 | 1920798 | 1922390 | + |
| XP_004511641.1 | LOC101488585 | Ca8 | 2152196 | 2153734 | + |
| XP_012574681.1 | LOC101509900 | Ca8 | 11064045 | 11068616 | + |
| XP_012574699.1 | LOC101501771 | Ca8 | 11070181 | 11077959 | + |
| XP_004512714.1 | LOC101510554 | Ca8 | 11084921 | 11090906 | + |
| XP_012574698.1 | LOC101503698 | Ca8 | 11196748 | 11199263 | - |
| XP_004513054.1 | LOC101489496 | Ca8 | 15856499 | 15857992 | + |
| XP_004513681.1 | LOC101501259 | Un | 15726 | 17333 | - |
| XP_004514511.1 | LOC101504040 | Un | 440047 | 451802 | - |
| XP_004515070.1 | LOC101494264 | Un | 14058 | 31344 | + |
| XP_027186553.1 | LOC113784536 | Un | 119199 | 119636 | - |
| XP_027186549.1 | LOC101507382 | Un | 138263 | 143089 | - |
| XP_004515891.1 | LOC101511223 | Un | 476974 | 478524 | + |
| XP_004516053.1 | LOC101500520 | Un | 2841 | 4310 | - |
| XP_004516070.1 | LOC101509930 | Un | 144146 | 148755 | + |

